# Immunosuppressive effect of the *Fusarium* secondary metabolite butenolide in human colon epithelial cells

**DOI:** 10.1101/2020.04.16.044388

**Authors:** Lydia Woelflingseder, Gerhard Adam, Doris Marko

## Abstract

Butenolide (BUT, 4-acetamido-4-hydroxy-2-butenoic acid gamma-lactone) is a secondary metabolite produced by several *Fusarium* species and is co-produced with the major trichothecene mycotoxin deoxynivalenol (DON) on cereal grains throughout the world. BUT has low acute toxicity and only very limited occurrence and exposure data are available. The intestinal epithelium represents the first physiological barrier against food contaminants. We aimed to elucidate the intestinal inflammatory response of the human, non-cancer epithelial HCEC-1CT cells to BUT and to characterize potential combinatory interactions with co-occurring trichothecenes, such as DON and NX-3. Using a reporter gene approach, BUT (≥5 *μ*M, 20 h) was found to decrease lipopolysaccharide (LPS; 10 ng/mL) induced nuclear factor kappa B (NF-κB) activation in a dose-dependent manner, and in combinatory treatments represses trichothecene-induced enhancement of this important inflammatory pathway. Analyzing transcription and secretion levels of NF-κB-dependent, pro-inflammatory cytokines, revealed a significant down-regulation of IL-1β, IL-6 and TNF-α in IL-1β-stimulated (25 ng/mL) HCEC-1CT cells after BUT exposure (10 *μ*M). Trichothecene-induced expression of pro-inflammatory cytokines by the presence of 1 *μ*M DON or NX-3 was substantially suppressed in the presence of 10 *μ*M BUT. The emerging mycotoxin BUT has the ability to suppress NF-κB-induced intestinal inflammatory response mechanisms and to modulate substantially the immune responsiveness of HCEC-1CT cells after trichothecene treatment. Our results suggest that BUT, present in naturally occurring mixtures of *Fusarium* fungal metabolites, should be increasingly monitored, and the mechanism of inhibition of NF-κB that might affect the pathogenesis or progression of intestinal inflammatory disorders, should be further investigated.

## INTRODUCTION

Mycotoxins are toxic secondary metabolites of low-molecular-weight produced by a wide diversity of filamentous fungi. Contaminating crops pre- or post-harvest, mycotoxins may enter the food and feed chains and pose a risk to human and animal health (Bennett and Klich, 2003). As mycotoxin-producing fungi, such as *Fusarium* species, are able to generate simultaneously a wide variety of secondary metabolites, co-exposure to mixtures of toxic fungal compounds is evident and the evaluation of potential toxicological interactions is therefore of great importance for comprehensive risk assessment.

Butenolide (BUT, 4-acetamido-4-hydroxy-2-butenoic acid gamma-lactone, Figure 1A) is a secondary metabolite produced by various *Fusarium* species, such as *F. sporotrichioides* and *F. graminearum.* It was already isolated from *F. tricinctum-*colonized tall fescue grass in the 1960s, causing a livestock mycotoxicosis called “fescue foot” (Yates et al., 1969; Grove et al., 1970; Tookey et al., 1972). Epidemiologically, BUT is considered one of the mycotoxins implicated in the etiopathogenesis of Kashin-Beck disease, an endemic, chronic and degenerative osteoarthropathy and of Keshan disease, an endemic cardiomyopathy, in China (Liu et al., 2007; Shi et al., 2009; Wang et al., 2009b). In regard to occurrence and exposure, scientific data are lacking. Abia et al. (2013) reported BUT in co-occurrence with trichothecene mycotoxins in up to 100% of the tested maize samples, resulting in mean contamination levels of 131 *μ*g/kg, respectively. A multi-mycotoxin analysis of Norwegian grain samples showed median BUT concentrations of about 200 *μ*g/kg, whereas the highest concentration of 3,370 *μ*g/kg was detected in oats (Uhlig et al., 2013).

**FIGURE 1:**
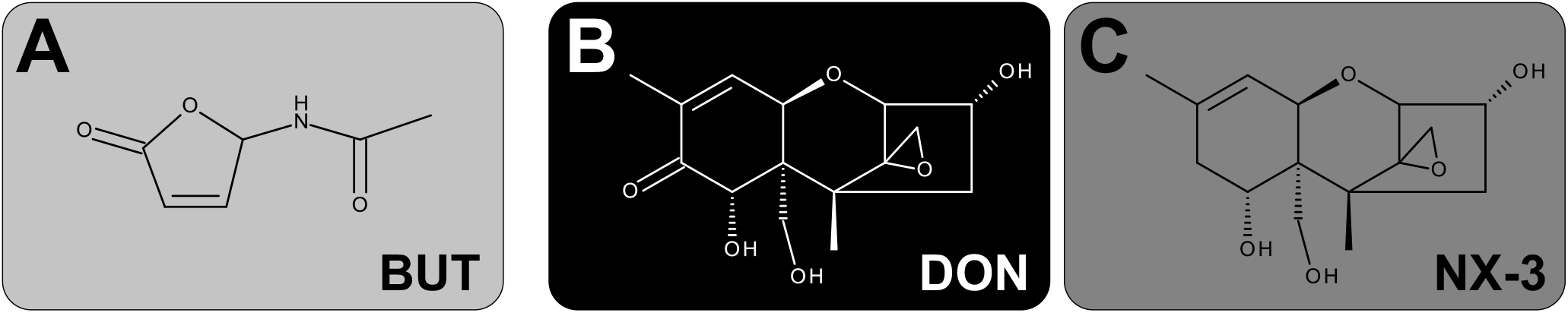
Chemical structures of the investigated secondary metabolites of Fusarium: (A) BUT, (B) DON and (C) NX-3.

After oral administration, BUT is of relatively low acute toxicity in mice (LD_50_ of 275 mg/kg body weight) (Yates et al., 1969), while in orally dosed steers 68 and 39 mg/kg caused death after 2 and 3 days, respectively (Tookey et al., 1972). Exposure to comparably high BUT concentrations (up to 1.4 mM) induced significant cytotoxic effects in various mammalian cell lines, mainly associated to its ability to cause oxidative damage and to trigger reactive oxygen species (ROS) production (Wang et al., 2006; Liu et al., 2007; Shi et al., 2009; Yang et al., 2010). In HepG2 cells, BUT concentrations in the low micromolar range where reported to substantially induce the redox-sensitive nuclear factor E2-related factor 2/antioxidant responsive element (Nrf2/ARE) pathway (Woelflingseder et al., 2018). Furthermore, the cellular glutathione pool was found to be crucial for cell survival after BUT treatment (Wang et al., 2006; Woelflingseder et al., 2018). Even though BUT is considered comparably not-toxic, recent toxicity studies assessing mixtures of BUT and deoxynivalenol (DON) revealed a significant contribution of BUT to the combined cytotoxicity in HepG2 cells (Woelflingseder et al., 2018). However, the impact of BUT on human intestinal epithelial cells and on their inflammatory response, as single compound and in mixtures, has not been investigated yet.

The *Fusarium* secondary metabolite DON (Figure 1B), also known as vomitoxin, is worldwide one of the most prevalent mycotoxins in temperate climate regions and therefore regarded as an important food safety issue (Kovalsky et al., 2016; EFSA et al., 2017). Its main mechanism of action is the inhibition of protein biosynthesis, triggered by an interaction of the epoxide moiety at C12-C13 with the peptidyltransferase at the 60S ribosomal subunit (Ueno, 1977; Garreau de Loubresse et al., 2014). The ribotoxic stress response leads to the activation of mitogen-activated protein kinases (MAPKs), which can be further related to aberrant gene regulation, cytokine production and induction of apoptosis (Iordanov et al., 1997; Laskin et al., 2002; Pestka et al., 2004). Several i*n vivo* studies reported DON to provoke intestinal inflammation (Azcona-Olivera et al., 1995; Zhou et al., 1997). In various human intestinal epithelial cancer cell lines, such as Caco-2, DON exposure (0.1-15 *μ*M; 48 h) caused a NF-κB dependent induction of interleukin-8 (IL-8) transcription and secretion, resulting in even stronger effects upon pro-inflammatroy stimulation by interleukin-1β(IL-1β), tumor necrosis factor-alpha (TNF-α) or lipopolysaccharide (LPS) (Moon et al., 2007; Maresca et al., 2008; Van De Walle et al., 2008). Regarding non-transformed, intestinal epithelial cells, IPEC-J2 porcine cells showed an upregulation of interleukin-1α (IL-1α), IL-1β, interleukin-6 (IL-6), IL-8, TNF-α and monocyte chemotactic protein 1 (MCP-1) after DON-exposure (2 *μ*M, 48 h), whereas at 0.5 *μ*M only IL-1β, IL-6 and IL-8 were induced (Wan et al., 2013). However, the modulatory effects of DON on the inflammatory response of human, non-cancer, colonic epithelial cells, especially focusing on the important inflammatory signaling pathway NF-κB, have not been assessed yet.

Whereas DON can be considered one of the most studied mycotoxins worldwide, only little is known about the recently discovered type A trichothecene NX-3 (Figure 1C). NX-3, structurally identified as an analogue of DON, lacking the carbonyl moiety at position C8, represents an *in planta* de-acetylation product of the *Fusarium* secondary metabolite NX-2, which is produced by an emerging population of *F. graminearum* in North America (Varga et al., 2015; Lofgren et al., 2018). A first toxicological characterization revealed NX-3 to inhibit *in vitro* plant- and mammalian-derived ribosomes to a similar extent as DON (Varga et al., 2015; Varga et al., 2018). Furthermore, both trichothecenes caused comparable cytotoxic effects in different mammalian cancer and non-cancer cell lines (Varga et al., 2018; Woelflingseder et al., 2018). Comparing the impact of NX-3 and DON on the onset of oxidative stress in HepG2 cells, only marginal differences were identified at the selected end-points (Woelflingseder et al., 2018). As DON has been reported to alter inflammatory signaling processes in human intestinal epithelial cells (Pestka et al., 2004), similar effects can be expected after exposure to the novel *Fusarium* mycotoxin NX-3. The assessment of specific effects in this regard might be crucial for hazard characterization and have not been assessed so far.

BUT, considered as moderately toxic, is rarely monitored and not regulated by any food safety authority, despite the fact that it can co-occur in high concentration with DON (Uhlig et al., 2013). Therefore we have assessed already in a previous study the ability of BUT to contribute to the toxicity of mixtures with trichothecene mycotoxins in HepG2 cells (Woelflingseder et al., 2018). So far, DON is well characterized with respect to its pro-inflammatory properties and several *in vitro* studies have already investigated the interaction of DON with other known mycotoxins on the immune response. However, a potential immunomodulatory contribution of less- or non-toxic co-occurring secondary metabolites, such as BUT, has not been taken into account so far and is the focus of the present study. In a first step, using a reporter gene approach, we assessed the impact of BUT, DON, NX-3 and respective combinations on the inflammatory key signaling pathway NF-κB. Furthermore, we analyzed mRNA expression and protein secretion levels of four pro-inflammatory cytokines performing qRT-PCR and Magnetic Luminex^®^ experiments, respectively, in order to obtain a more detailed characterization of the inflammatory response of the applied non-cancer intestinal cell model HCEC-1CT.

## MATERIALS AND METHODS

### Chemicals and reagents

BUT was synthesized and purified according to Burkhardt et al. (1968), as described previously by Woelflingseder et al. (2018). DON was purchased from Romer Labs (Tulln, Austria). NX-3 was produced and purified as deacetylation product of NX-2 using preparative high-performance liquid chromatography (purity > 99% according to LC-UV at 200 nm) as published by Varga et al. (2015). All substances were dissolved in water (LC-MS grade) to obtain stock solutions of 10 mM, which were aliquoted and stored at −20 °C. Cell culture media, supplements and material were purchased from GIBCO Invitrogen (Karlsruhe, Germany), Sigma-Aldrich (Munich, Germany) and Sarstedt AG & Co (Nuembrecht, Germany).

### Cell culture and treatment

The human monocytic cell line THP1-Lucia™ NF-κB, deriving from the human THP-1 monocyte cell line by stable integration of an NF-κB inducible Luciferase reporter construct, was purchased from InvivoGen (San Diego, USA). The non-tumorigenic human colon epithelial cell line HCEC-1CT (Roig et al., 2010; Roig and Shay, 2010) was kindly provided by Prof. Jerry W. Shay (UT South-western Medical Center, Dallas, TX, USA). THP1-Lucia™ were cultured in RPMI 1640 medium, supplemented with heat-inactivated 10% (v/v) fetal bovine serum and 1% (v/v) penicillin/streptomycin (100 U/mL). THP1-Lucia™ were treated alternately with zeocin and normocin (100 *μ*g/mL; Invivogen, USA). HCEC-1CT cells were cultivated in Dulbecco’s Modified Eagle’s Medium (high glucose) supplemented with 2% (v/v) Medium 199 (10X), 2% (v/v) GE Healthcare™ HyClone™ Cosmic Calf™ Serum, 20 mM 4-(2-hydroxyethyl)-1-piperazineethanesulfonic acid, 50 *μ*g/mL gentamicin, insulin-transferrin-selenium-G (10 *μ*g/mL; 5.5 *μ*g/mL; 6.7 ng/mL), recombinant human epidermal growth factor (18.7 ng/mL) and hydrocortisone (1 *μ*g/mL). Both cell lines were maintained in humidified incubators at 37°C and 5% CO_2_ and sub-cultured every 3-4 d. Cells were routinely tested for the absence of mycoplasm contamination and used for experiments at passages 12-18. BUT, DON, NX-3 and their combinations were added to the incubation solutions, resulting in a final solvent concentration of 2% (v/v) water (LC-MS grade). In order to ensure data comparability, combinatory treatments were always performed in parallel to the individual substances using the same passage of cells on the same microtiter-plate.

### NF-κB reporter gene assay in THP1-Lucia™ cells

BUT, DON, NX-3 and 10:1 and 1:1 combinations of BUT with the trichothecenes were prepared in reaction tubes to be further diluted 1:100 by the addition of the THP1-Lucia™ cell suspension. 10:1 and 1:1 ratios were selected due to the high variability of secondary metabolite concentrations in the available occurrence studies (Abia et al., 2013; Streit et al., 2013a; Uhlig et al., 2013) and due to previously performed cell viability experiments in Woelflingseder et al. (2018). Cellular concentration was determined by trypan blue exclusion, cells were centrifuged for 5 min at 250 x g and resuspended at a concentration of 1 x 10^6^ cells/mL in fresh, pre-warmed growth medium. Cell suspension was added to the respective toxin preparations into the reaction tubes. They were gently mixed and transferred as technical duplicates into a 96-well plate (100 *μ*L/well). After 2 h of incubation at 37°C and 5% CO_2_ in the humidified incubator, LPS (1 *μ*L/well) at a final concentration of 10 ng/mL was added to the cells and incubated further for 18 h. 2% (v/v) water (LC-MS grade) with and without LPS treatment were used as solvent control. As recommended by the supplier, heat killed *Listeria monocytogenes* (HKLM) at a concentration of 20 x 10^6^ cells/well were applied as positive control for TLR activity. Following in total 20 h of incubation, well plate was centrifuged (250 x g, 5 min) and 10 *μ*L/well of the supernatant were collected. Subsequently, the reporter gene activity assay (secreted luciferase) was performed according to the manufacturer’s protocol. QUANTI-Luc™ solution (InvivoGen, US), containing the recommended coelenterazine substrate, was added for luminescence measurements.

In parallel cellular metabolic activity was monitored by the alamarBlue^®^ assay (for details see “Determination of cellular metabolic activity and protein content”). Therefore, after supernatant collection for the measurements of luciferase activity, 10 *μ*L of alamarBlue^®^ reagent (Thermo Fisher Scientific, Massachusetts, USA) were added to the cells and incubated for 2 h. Subsequently, after another centrifugation of the plate (250 x g, 5 min) 70 *μ*L/well were transferred to a black 96-well plate and fluorescence intensity was measured at 530/560 nm (excitation/emission). Both luciferase activity and fluorescence intensity measurements were performed on a SynergyTM H1 hybrid multi-mode reader (BioTek, Bad Friedrichshall, Germany) assessing at least five independent experiments in technical duplicates.

### Quantitative analysis of cytokine gene transcription in HCEC-1CT cells

In HCEC-1CT cells gene transcription of the four cytokines TNF-α, IL-1β, IL-8 and IL-6 was analyzed by quantitative reverse transcription polymerase chain reaction (qRT-PCR). Cells were seeded in 12-well plates (35,000 cells/well) and allowed to grow for 48 h. Cells were incubated with BUT, DON, NX-3 and respective combinations in total for 5 h, consisting of 2 h of toxin incubation, followed by IL-1β co-treatment (25 ng/mL) for additional 3 h. Subsequently, total RNA was extracted using Maxwell® 16 Cell LEV Total RNA Purification Kits (Promega, US) and reversed transcribed into complementary DNA (cDNA) by QuantiTect^®^ Reverse Transcription Kit (Qiagen) according to the supplier’s protocols. Afterwards, cDNA samples were amplified in technical duplicates in presence of gene specific primers (QuantiTect^®^ Primer Assays, Qiagen) and QuantiTect^®^ SYBR Green Master Mix (Qiagen) using a StepOnePlus™ System (Applied Biosystems, Foster City, USA). Therefore, the following primer assays were used: β-actin (ACTB1, Hs_ACTB1_1_SG, QT00095431); glyceraldehyde 3-phosphate dehydrogenase (GAPDH, Hs_GAPDH_1_SG, QT000079247); TNF-α (Hs_TNF_1_SG; QT00029162); IL-1β (Hs_IL1B_1_SG, QT00021385); IL-8 (Hs_CXCL8_1_SG, QT00000322); IL-6 (Hs_IL6_1_SG, QT00083720). A universal PCR protocol was applied including enzyme activation at 95 °C for 15 min, 40 cycles of 15 s at 94 °C, 30 s at 55 °C and 30 s at 72 °C. In order to check for primer specificity melting curve analysis was performed at the end of every PCR experiment. StepOnePlus^®^ software (version 2.3, Applied Biosystems, USA) was used for fluorescence signal quantification and further data analysis. Of each tested sample at least five independent experiments were performed. Presented transcript data was normalized to the mean of transcript levels of endogenous control genes (ACTB1, GAPDH) applying the ΔΔCt-method (Schmittgen and Livak, 2008) for relative quantification.

### Profiling of pro-inflammatory cytokine secretion

Cytokines released from HCEC-1CT cells into cell culture supernatants after 5 h of incubation were measured using a Magnetic Luminex^®^ assay (R&D SYSTEMS, Minneapolis, USA). Cell culture supernatant samples were collected after the incubation for cytokine gene transcription (details for incubation see 2.4.) and stored until the measurement at −80 °C. After 2-fold sample dilution for TNF-α analysis and 40-fold dilution for the analysis of IL-8 and IL-6, the assay was performed according to the manufacturer’s protocol. Briefly, in a 96-well plate 50 *μ*L of standard or sample were mixed with 50 *μ*L microparticle cocktail per well. After 2 h of incubation at room temperature (RT), plate was washed three times with 100 *μ*L wash buffer per well. Afterwards, samples were incubated with 50 *μ*L per well of a biotin-antibody cocktail for 1 h at RT. Again, three washing steps were performed prior to 30 minutes incubation with 50 *μ*L per well of streptavidin-PE solution. Finally, samples were rinsed again three times with wash buffer and resuspended in 100 *μ*L per well prior to the read-out on a Bio-Plex^®^ 200 multiplex array reader (Bio-Rad Laboratories, Hercules, USA). Using the Bio-Plex Manager™ software a standard curve was created for each analyte by performing a five-parameter logistic curve-fit, average blank-values were subtracted and respective cytokine concentrations were calculated. The assay was performed in technical duplicates, analyzing at least three independent biological replicates.

### Determination of cellular metabolic activity and protein content (alamarBlue® and sulforhodamine B assay)

In order to rule out combinatory effects and cytotoxicity on gene transcription analysis, the impact of BUT, DON, NX-3 and their combinations on cell viability after 5 h of incubation was determined. Therefore, two cell viability assays, the alamarBlue^®^ and the sulforhodamine B assay (SRB) according to Skehan et al. (1990) were performed. HCEC-1CT cells were seeded into 96-well plates (2,500 cells/well) and allowed to grow for 48 h. Cells were then incubated for a total of 5 h, consisting of 2 h of toxin pre-incubation, followed by co-treatment with 25 ng/mL IL-1β for 3 h. As BUT was combined in a constant 10:1 ratio with DON and NX-3, the following concentrations and combinations were applied, obtained by 1:100 dilutions of respective stock solutions: 0.1-100 *μ*M for BUT; 0.01-10 *μ*M for DON/NX-3. After 4 hours of incubation, 10 *μ*L alamar-Blue^®^ reagent (Thermo Fisher Scientific, Massachusetts, USA) were added to the cells and incubated for further 1.25 h. Subsequently, 70 *μ*L/well were transferred to a black 96-well plate and fluorescence intensity was measured at 530/560 nm (excitation/emission) using a SynergyTM H1 hybrid multi-mode reader (BioTek, Bad Friedrichshall, Germany). Afterwards, cells were rinsed with pre-warmed phosphate buffered saline and fixed by the addition of 10 *μ*L/well 5% (v/v) trichloroacetic acid. After 30 min of incubation at 4 °C, wells were washed three times with water and dried overnight at RT. Cells were incubated for 1 h with 0.4% (w/v) SRB in 1% (v/v) acetic acid and then washed twice with water and with 1% (v/v) acetic acid solution to remove remaining staining solution. Plates were dried in the dark at RT. Before reading single wavelength absorbance at 570 nm with the SynergyTM H1 hybrid multi-mode reader, dye was dissolved in 100 *μ*L/well 10 mM Tris buffer (pH 10). A solvent control with and without IL-1β treatment were used. Cell-free blank values were subtracted and measured data of at least five independent, biological replicates performed in technical duplicates, was referred to the IL-1β-stimulated solvent control.

### Data visualization and statistical analysis

For statistical analysis and data visualization the software Origin 2018 (Northampton, USA) was used. In the reporter gene and cell viability experiments respective data were tested for normality of the distribution with the Shapiro-Wilk test. Significant differences between concentrations of the same compound with the respective lowest tested concentration were calculated by one‐way ANOVA, followed by Bonferroni’s post-hoc tests (*p* < 0.05), whereas significant differences between the DON or NX-3 and the respective combinations with BUT were calculated with Student’s *t-*test. Regarding the gene transcription and cytokine secretion data, significant differences were calculated with Student’s *t-*test.

## RESULTS

### Impact on NF-κB signaling pathway activity

In order to elucidate the role of BUT, DON, NX-3 and their combinations on NF-κB signaling pathway activation, THP1-Lucia™ cells, carrying a NF-κB-inducible Luc reporter construct, were grown 20 h in the presence of respective *Fusarium* secondary metabolites. After 2 h incubation, cells were further exposed to a pro-inflammatory IL-1β-stimulus (25 ng/mL). In a first experiment equimolar concentrations of BUT and the trichothecenes over a concentration range of 0.01 to 10 *μ*M were tested (Figure 2A).

**FIGURE 2:**
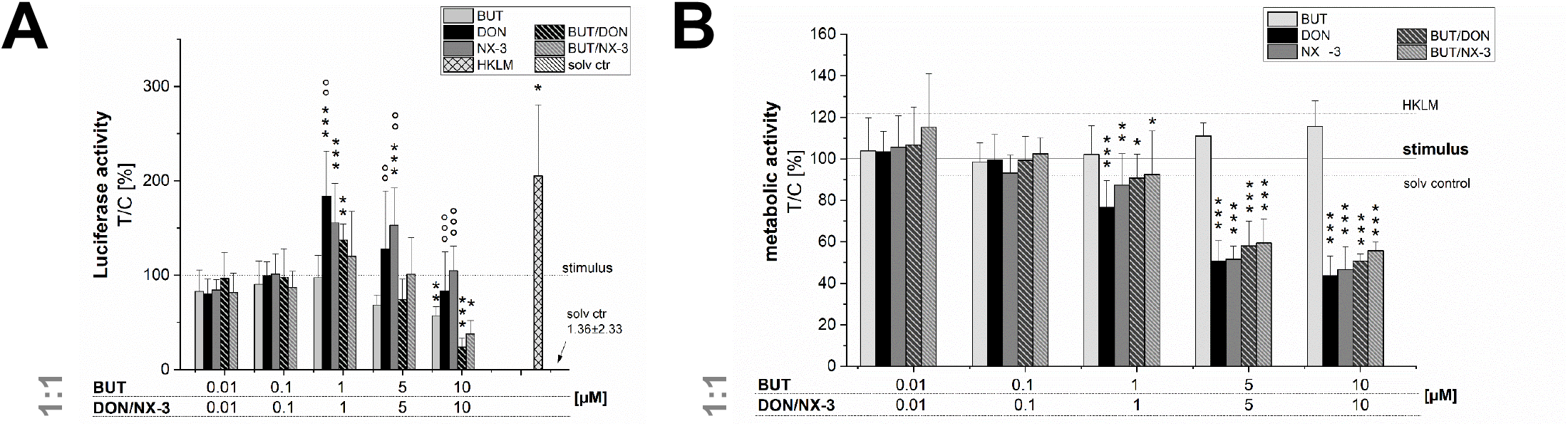
Effect of toxin pre-treatment on NF-κB-dependent luciferase (ratio LPS treated/untreated control). Modulation of NF-κB-dependent luciferase expression (A) after 2 h pre-incubation with BUT (0.01–10 *μ*M), DON (0.01-10 *μ*M), NX-3 (0.01-10 *μ*M) and respective equimolar combinations, followed by 18 h co-stimulation with LPS (10 ng/mL) in THP1-Lucia™ cells referred to the respective alamarBlue^®^-fluorescence intensity (B). 2% (v/v) water (LC-MS grade) with (dotted line) and without LPS treatment (solv ctr) served as solvent control, HKLM (20 x 10^6^ cells/well) as positive control. alamarBlue^®^-fluorescence intensity and luciferase activity data are expressed as mean values ± SE of at least five independent experiments performed in duplicates normalized to the LPS-stimulated solvent control (2% water (LC–MS grade)). Significant differences to the stimulated solvent control are indicated with * (p < 0.05), ** (p < 0.01) and *** (p < 0.001), whereas significant differences of the combinatory data to the respective DON and NX-3 data are indicated with ° (p < 0.05), °° (p < 0.01) and °°° (p < 0.001).

The data show the percentage change of reporter gene activity of treatment (LPS + mycotoxin) in relation to LPS treatment alone. Compared to the solvent control already LPS alone triggered a 270-fold increase in luciferase activity, which was further enhanced by the co-treatment with the trichothecenes. Serving as positive control, heat killed *Listeria monocytogenes* cells (HKLM) induced the reporter stronger than LPS alone. Whereas the lowest tested BUT concentrations (0.01, 0.1 and 1 *μ*M) did not modulate NF-κB activity, at 5 *μ*M a reduced luciferase signal was detected, reaching statistical significance at 10 *μ*M. Both trichothecenes, DON and NX-3 caused similar effects, so that at 1 *μ*M a significant induction of the luciferase signal up to 183 ± 48% (DON) and 155 ± 42% (NX-3) of the stimulated solvent control were observed. These inductive effects were still measurable with 5 *μ*M DON or NX-3. However, at 10 *μ*M of either toxin a decrease of the signal to the solvent control level was determined, in line with a substantial loss in metabolic activity seen in the alamarBlue^®^ assay (Figure 2B). In comparison to the effects induced by DON or NX-3 as single compounds, equimolar combinations of BUT and the trichothecenes at concentrations ≥ 1 *μ*M, resulted mostly in a lower luciferase signal.

Thus, on the one hand 1 *μ*M DON and NX-3 led to a potent induction of the NF-κB signaling pathway and on the other hand 10 *μ*M BUT significantly reduced the signal to 57 ± 10% without any cytotoxicity involved. Therefore, further experiments were performed combining BUT at a 10-fold molar excess of the trichothecene mycotoxins (Figure 3). 50 and 100 *μ*M BUT, only the latter causing a significant decrease in cell viability, reduced the NF-κB activity to 0.3 ± 0.2% and 0.1 ± 0.1%, respectively. In 10:1 combination with 1 *μ*M DON or NX-3, the presence of BUT reduced the luciferase signal from 195% and 196% to a signal even below the control level (96% and 91%). At higher concentrations, cytotoxic effects need to be taken into account. Interestingly, NF-κB activity was completely suppressed in mixtures containing 5 and 10 *μ*M DON or NX-3.

**FIGURE 3:**
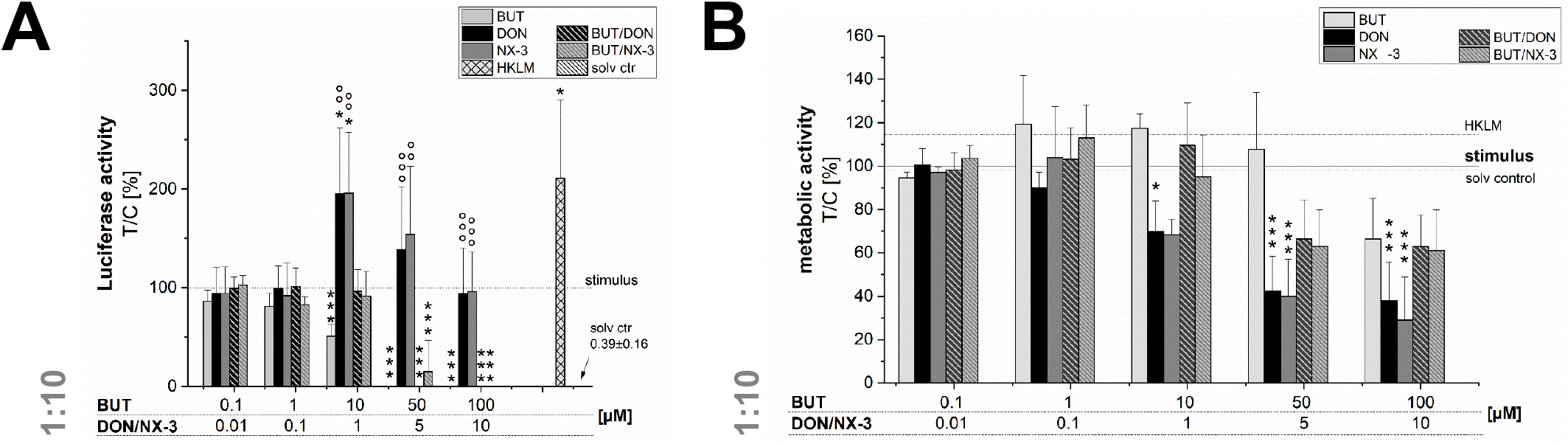
Modulation of NF-κB-dependent luciferase expression (A) after 2 h pre-incubation with BUT (0.1–100 *μ*M), DON (0.01-10 *μ*M), NX-3 (0.01-10 *μ*M) and respective 10:1 combinations, followed by 18 h co-stimulation with LPS (10 ng/mL) in THP1-Lucia™ cells referred to the respective alamarBlue^®^-fluorescence intensity (B). 2% (v/v) water (LC-MS grade) with (dotted line) and without LPS treatment (solv ctr) served as solvent control, HKLM (20 x 10^6^ cells/well) as positive control. alamarBlue^®^-fluorescence intensity and luciferase activity data are expressed as mean values ± SE of at least five independent experiments performed in duplicates normalized to the LPS-stimulated solvent control (2% water (LC–MS grade)). Significant differences to the stimulated solvent control are indicated with * (p < 0.05) and *** (p < 0.001), whereas significant differences of the combinatory data to the respective DON and NX-3 data are indicated with °° (p < 0.01) and °°° (p < 0.001).

### Modulatory effects of BUT on trichothecene-induced cytokine gene transcription

NF-κB signaling is involved in transcriptional induction of many cytokines, chemokines and additional inflammatory mediators, such as interleukins, lymphokines and tumor necrosis factors (Hayden and Ghosh, 2011; Sun et al., 2013; Liu et al., 2017). As BUT was found to suppress significantly LPS-induced NF-κB activation in the THP1-Lucia™ cell line and to further inhibit NF-κB induction after trichothecene exposure, we aimed to elucidate the impact of BUT at the level of gene transcription of four NF-κB-regulated genes (TNF-α, IL-1β, IL-8 and IL-6) in HCEC-1CT cells (Figure 4).

**FIGURE 4:**
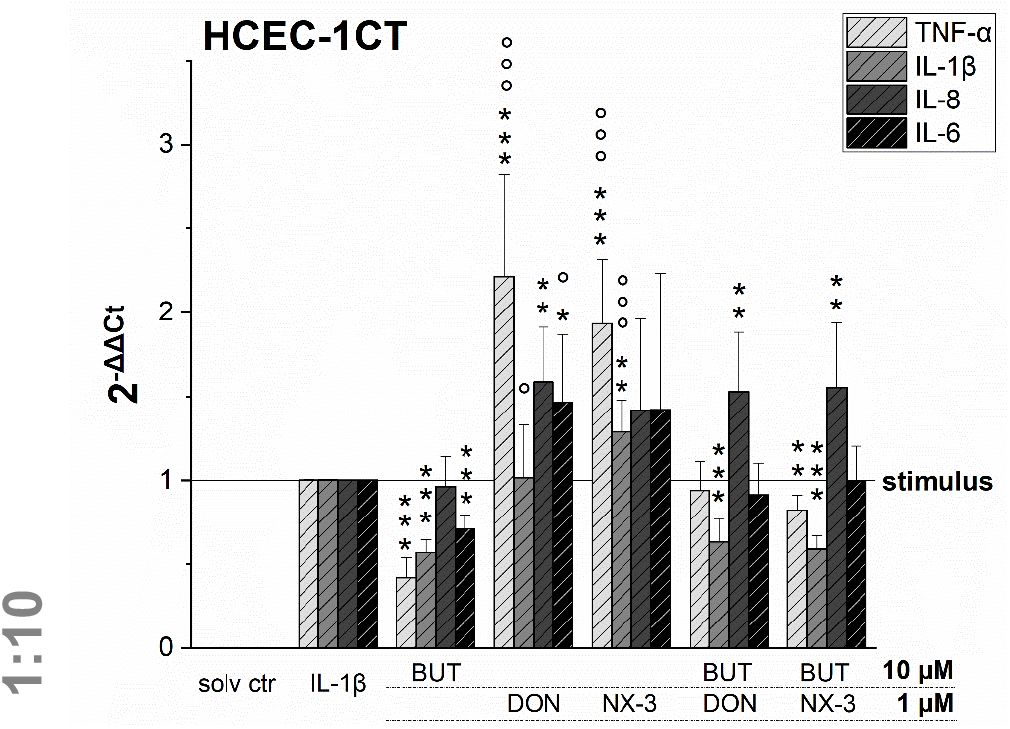
Gene transcription levels of TNF-α, IL-1β, IL-8 and IL-6 in HCEC-1CT cells after 2 h pre-incubation with BUT (10 *μ*M), DON (1 *μ*M), NX-3 (1 *μ*M) and respective 10:1 combinations, followed by 3 h co-stimulation with IL-1β (25 ng/mL) measured by qRT-PCR. Transcription data are normalized to the gene expression levels of two endogenous control genes (ACTB1 and GAPDH) and referred to the stimulated solvent control levels. Data are expressed as mean values ± SD of at least six independent experiments performed in technical duplicates. Significant differences to the stimulated solvent control are indicated with ** (p < 0.01) and *** (p < 0.001), whereas significant differences of the combinatory data to the respective DON and NX-3 data are indicated with ° (p < 0.05) and °°° (p < 0.001).

Cells pre-exposed for 2 h to BUT (10 *μ*M) and then further challenged for 3 h with IL-1β (25 ng/mL), showed a significantly reduced mRNA expression of TNF-α, IL-1β and IL-6. Whereas IL-8 was not altered significantly in its gene transcription compared to the IL-1β stimulated solvent control samples, only 0.4-, 0.6- and 0.7-fold expressions of TNF-α, IL-1β and IL-6 transcripts were reported, respectively. DON and NX-3 (1 *μ*M), known for their pro-inflammatory properties, caused a substantial increase in TNF-α, IL-8 and IL-6 mRNA expression, while IL-1β gene transcription was found to be significantly elevated only after NX-3 treatment. Finally, combinations of BUT (10 *μ*M) with DON or NX-3 (both 1 *μ*M) were analyzed in the qRT-PCR experiments. For both trichothecenes a similar pattern of cytokine mRNA levels was determined. TNF-α, induced after DON and NX-3 treatment 2.2- and 1.9-fold, was reduced to 0.9 and 0.8-fold relative transcription, respectively, in combinatory treatments with BUT. While a similar reduction to the IL-1β stimulated solvent control level was found for IL-6, an even stronger decrease was determined in case of IL-1β, which was reduced to relative transcription levels of approximately 0.6. However, once more, IL-8 transcription levels were not affected by BUT in the combinatory treatments.

### Impact of BUT and its combinations with DON and NX-3 on cytokine secretion

Quantifying cytokine secretion levels by a magnetic Luminex® assay, we aimed to investigate, whether the reduced cytokine gene transcription of TNF-α, IL-8 and IL-6 after BUT-treatment, is also reflected on the protein level (Figure 5). 10 *μ*M BUT significantly reduced TNF-α and IL-6 expression levels to 62 and 81% of the IL-1β stimulated solvent control samples, respectively. Protein levels of IL-8 were slightly induced after BUT treatment. Trichothecene treatments (1 *μ*M) did not cause significant alterations of the cytokine secretion levels of TNF-α, IL-8 and IL-6 compared to the IL-1β stimulated control levels. However, combinatory treatments of BUT with DON or NX-3 showed a similar pattern as already observed after BUT treatment: TNF-α and IL-6 expression levels were both significantly reduced to 67% in case of DON and to 69 and 68%, respectively, after NX-3-exposure. In regard to IL-8 secretion also in the combinatory treatments no significant modulations were observed. Comparing the different cytokine concentrations after IL-1β stimulation (Table 1) substantial differences were found. Whereas TNF-α was present at very low concentrations of about 50 pg/mL, IL-6 was detected at concentrations around 2,700 pg/mL. IL-8 was the most abundant cytokine, occurring at concentrations around 46 ng/mL.

**FIGURE 5:**
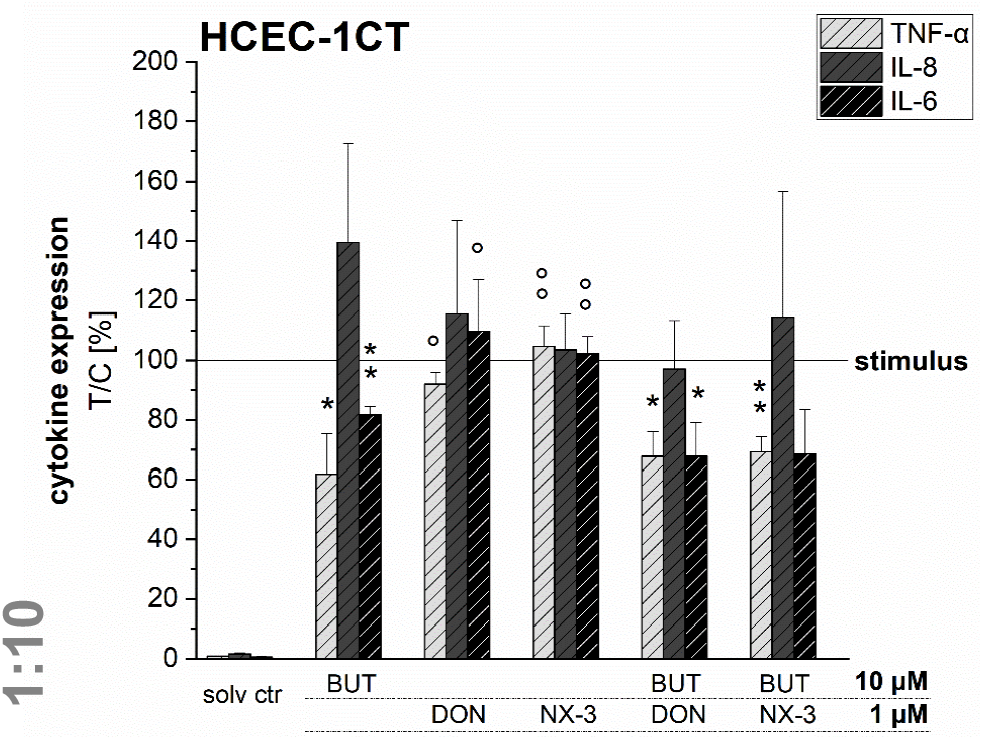
Cytokine expression levels of TNF-α, IL-8 and IL-6 in HCEC-1CT cells after 2 h pre-incubation with BUT (10 *μ*M), DON (1 *μ*M), NX-3 (1 *μ*M) and respective 10:1 combinations, followed by 3 h co-stimulation with IL-1β (25 ng/mL) measured by a Magnetic Luminex^®^ assay. Cytokine expression data are normalized to the stimulated solvent control levels. Data are expressed as mean values ± SD of at least three independent experiments performed in technical duplicates. Significant differences to the stimulated solvent control are indicated with * (p < 0.05) and ** (p < 0.01), whereas significant differences of the combinatory data to the respective DON and NX-3 data are indicated with ° (p < 0.05) and °° (p < 0.01).

**TABLE 1:**
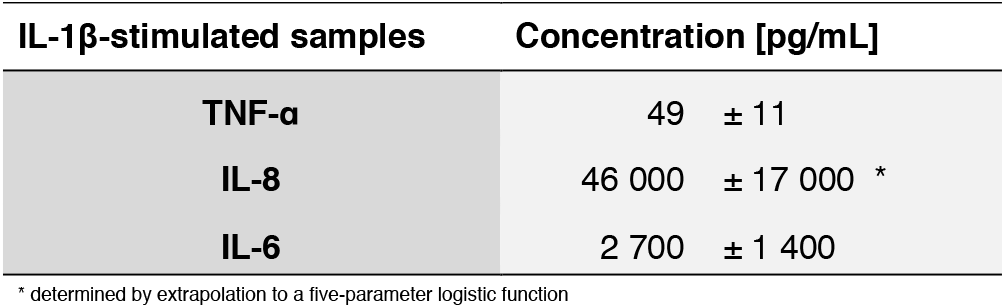
Absolute cytokine expression levels in pg/mL of TNF-α, IL-8 and IL-6 of HCEC-1CT measured in the supernatants of the IL-1β-stimulated solvent control samples by a Magnetic Luminex^®^ assay.

| IL-1β-stimulated samples | Concentration [pg/mL] |
| --- | --- |
| TNF-α | 49 ± 11 |
| IL-8 | 46 000 ± 17 000 * |
| IL-6 | 2 700 ± 1 400 |

\* determined by extrapolation to a five-parameter logistic function

### Assessment of cell viability after exposure to BUT and its combinations with DON and NX-3

In order to rule out potential cytotoxicity to interfere in the qRT-PCR experiments, cell viability was assessed after 5 h exposure. In the alamarBlue^®^ assay cellular metabolic activity was elucidated, whereas in parallel the cellular protein content was determined in the SRB assay (Figure 6). BUT decreased significantly the metabolic activity at 50 and 100 *μ*M after 2 h pre-incubation and 3 h co-exposure with IL-1β. However, in the SRB assay no significant effects of BUT on the protein content of HCEC-1CT cells were determined.

**FIGURE 6:**
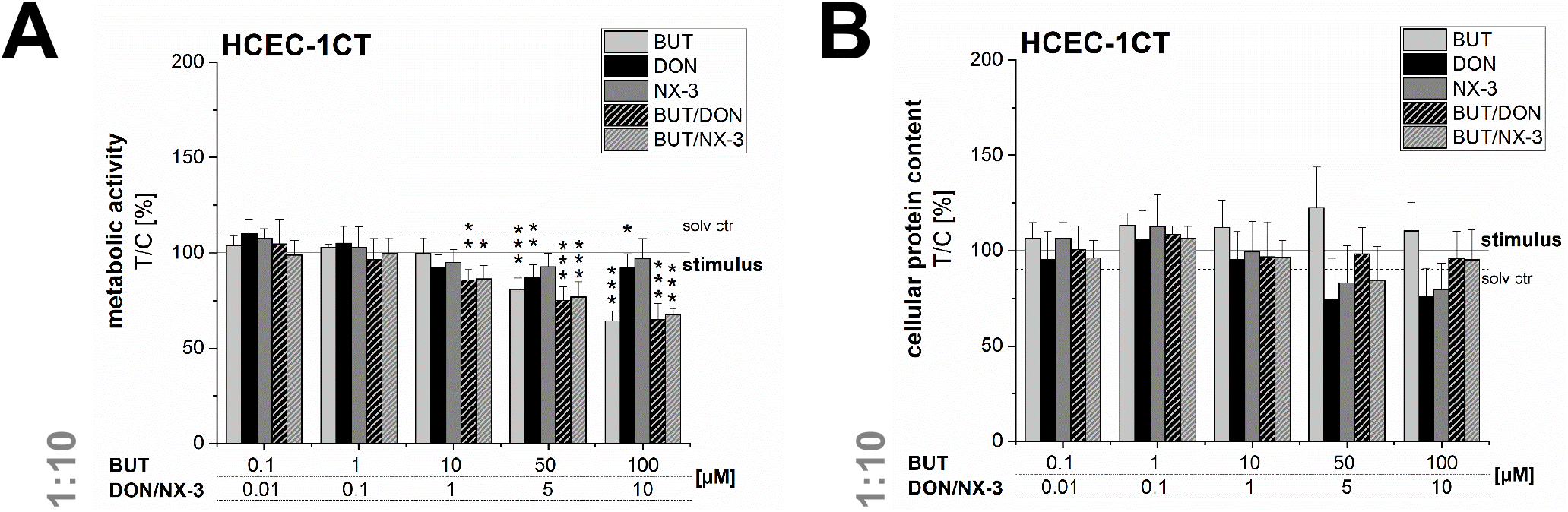
Effects on cell viability of HCEC-1CT cells after 2 h pre-incubation with BUT (10 *μ*M), DON (1 *μ*M), NX-3 (1 *μ*M) and respective 10:1 combinations, followed by 3 h co-stimulation with IL-1β (25 ng/mL) in the alamarBlue^®^ (A) and the SRB-assay (B). Cell viability data are normalized to the stimulated solvent control levels (solid line, dotted line: solvent control without pro-inflammatory stimulus). Data are expressed as mean values ± SD of at least five independent experiments performed in technical duplicates. Significant differences to the stimulated solvent control are indicated with * (p < 0.05), ** (p < 0.01) and *** (p < 0.001).

At the highest concentrations tested (5 and 10 *μ*M), DON and NX-3 slightly reduced the fluorescence signal in the alamarBlue^®^ assay, even though NX-3 showed the tendency to be less potent. However, these effects were found to be more pronounced in the SRB assay reducing the cellular protein content in case of DON to 74 ± 21% (5 *μ*M) and 76 ± 15% (10 *μ*M) of the IL-1β challenged solvent control and to 83 ± 20% (5 *μ*M) and 80 ± 1% (10 *μ*M) after NX-3 incubation.

Mixtures of BUT with the trichothecenes at a constant molar ratio of 10:1, caused a slight loss in metabolic activity at 10/1 *μ*M. Higher concentrations (50/5 and 100/10 *μ*M) resulted in a more pronounced effect, reducing the metabolic activity to about 76 and 66%. However, in all tested combinations the impact of DON and NX-3 was found similar. Regarding the cellular protein content, up to the highest tested concentrations only marginal effects were observed in the SRB assay.

## DISCUSSION

As the intestinal epithelium represents an important defense barrier against food contaminants, numerous studies investigated the impact of *Fusarium* mycotoxins on intestinal cells and mediated immune and inflammatory response mechanisms (Van De Walle et al., 2008; Wan et al., 2013; Pinton and Oswald, 2014; Alassane-Kpembi et al., 2017). Although the most prevalent *Fusarium* mycotoxin DON often co-occurs with many other co-regulated secondary fungal metabolites, data describing potential combinatory effects on intestinal inflammation are lacking. Especially fungal compounds of low acute toxicity (Yates, 1971; Tookey et al., 1972), however occurring at high concentrations, such as BUT (Abia et al., 2013; Streit et al., 2013b; Uhlig et al., 2013; Ivanova et al., 2016), are rarely taken into account.

To the best of our knowledge, so far effects of BUT on the activation of the NF-κB signaling pathway in intestinal epithelial cells have not been reported. The transcription factor NF-κB is considered one of the key regulators of inflammatory immune responses and its activation alters the expression of several inflammatory mediators, including cytokines and enzymes like cyclooxygenase-2 (COX-2) or inducible nitric oxide synthase (iNOS) (Liu et al., 2017). Using a reporter gene approach, we elucidated, applying not-cytotoxic BUT concentrations (≤ 50 *μ*M), significant reductions in NF-κB activity (Figure 2 and Figure 3). Previously, various butenolide-based structures, belonging to the family of α, β-unsaturated lactones, known as furanones, but decorated with different moieties, partly showed anti-inflammatory activity (Khokra et al., 2013; Ali et al., 2015; Liu et al., 2018). While Ali et al. (2015) reported suppressive effects of novel butenolide-based benzyl pyrrolones (20 *μ*M) on TNF-α, COX-2 and NF-κB protein expression of RAW 264.7 cells after 24 h incubation, *in silico* docking studies revealed other butenolide-derivatives as potential COX-2 inhibitors (Khokra et al., 2013). Recently anti-inflammatory capacities of ten butenolide-derivatives, isolated from the coral-derived fungus *Aspergillus terreus* were elucidated. Three compounds, tested at a concentration of 20 mM, showed remarkable inhibitory effects against nitric oxide production in the LPS-stimulated murine macrophage cell line, RAW 264.7 (Liu et al., 2018).

Even though preliminary studies described anti-inflammatory properties of various butenolide-derivatives, little is known about the underlying mechanism of action. However, as the *Fusarium* secondary metabolite BUT was found to significantly activate the Nrf2/ARE signaling pathway at concentrations between 1 and 50 *μ*M (Woelflingseder et al., 2018) and as Nrf2 and NF-κB are considered two interplaying key pathways regulating the cellular redox status and responses to inflammatory triggers, their molecular cross-talk might play an important role in BUT-induced toxicities. One possible hypothesis to be tested in future work is that BUT, as a Michael acceptor, potentially reacts with the critical cysteine thiolate groups of the Nrf2-repressor protein Kelch-like ECH-associated protein 1, releasing Nrf2 and inducing the transcription of ARE-mediated antioxidative and cytoprotective enzymes, such as heme oxygenase-1 (HO-1), NAD(P)H dehydrogenase (quinone 1), γ-glutamate-cysteine ligase, glutathione peroxidase and glutathione S-transferases (Luo et al., 2007; Nakamura and Miyoshi, 2010; Magesh et al., 2012). Numerous studies dealing with Nrf2-activating compounds reported respective anti-inflammatory potential, including reduced COX-2, TNF-α, iNOS and IL-1β production (Lin et al., 2008; Wardyn et al., 2015). Furthermore, increases in glutathione levels, as well as in HO-1 activity were found to decrease or to even inhibit NF-κB activity (Soares et al., 2004; Jiang et al., 2014; Wardyn et al., 2015). Hence, it can be speculated that BUT inhibits NF-κB activity in a negative feedback loop due to a pronounced Nrf2/ARE signaling pathway activation.

Combining BUT, as an NF-κB-suppressor, at equimolar concentrations or 10-fold excess of BUT with the co-occurring mycotoxins DON and NX-3, both causing an increase in NF-κB activity, mediated a reduction of the luciferase signal down to solvent control level (Figure 2 and Figure 3). Whereas the impact of NX-3 on the activation of the NF-κB signaling pathway has not been addressed in literature so far, DON-induced NF-κB translocation has been reported already several times (Katika et al., 2015; Adesso et al., 2017). This trichothecene-induced inflammatory response was further enhanced by co-treatment with LPS and interferon-γ of non-tumorigenic rat IEC-6 cells, involving TNF-α production, iNOS and COX-2 expression, ROS release and inflammasome activation (Adesso et al., 2017). In intestinal explants and human Caco-2 cells an induction of gene expression and secretion of the NF-κB-dependent, pro-inflammatory cytokine IL-8 was observed after exposure to DON (Van De Walle et al., 2008; Alassane-Kpembi et al., 2017). This interleukin is considered an early marker of inflammatory processes, acting as a potent chemo-attractant for leukocytes and T-lymphocytes underlying gut epithelial cells (Warhurst et al., 1998). In our study both trichothecenes (1 *μ*M) increased mRNA levels of IL-8, as well as of TNF-α and IL-6 (Figure 4). BUT, in contrast, significantly reduced transcript concentrations of TNF-α, IL-1β and IL-6, whereas no suppression was found in case of IL-8. Accordingly, in the combinatory cell treatments, a suppression of TNF-α, IL-6 and IL-1β transcripts was found, of which the latter was even significantly below the stimulated solvent control level. IL-8 was not modified by the presence of BUT, showing similar mRNA levels as in the trichothecene-treated samples. Going one step further, having a look at potential modifications of the protein expression levels (Figure 5), surprisingly the induction in the transcription levels of all four pro-inflammatory cytokines due to the presence of DON and NX-3 was not reflected in the cytokine expression. Van De Walle et al. (2008), applying similar DON concentrations (1 and 10 *μ*M) on Caco-2 cells for 48 h in presence of IL-1β (25 ng/mL), reported a dose-dependent increase of the IL-8 secretion levels, in accordance with the findings of Moon et al. (2007) in human intestinal 407 cells after 12 h incubation. Corresponding to the substantial inhibition of the NF-κB signaling pathway observed in the reporter gene assay, BUT, alone, as well as in the combinatory treatments significantly reduced TNF-α and IL-6 cytokine expression. However, it remains unclear, why BUT has no inhibitory effect on IL-8 transcription and expression, even though a substantial suppression of the NF-κB activity leading to a decrease in TNF-α, IL-6 and IL-1β transcription and secretion was detected. Cytokine secretion is multifactorially regulated. Having a closer look into the IL-1 signaling pathway, IL-1β-stimulation causes not only post-receptor activation of NF-κB, but also of MAPKs such as p38 or c-Jun N-terminal kinases/stress-activated kinases (JNK/SAPK), cooperatively inducing the transcription of target genes like IL-6, IL-1β and IL-8 (Dunne and O’Neill, 2003; Weber et al., 2010). Furthermore, the differences in the cytokine concentrations determined in the Magnetic Luminex® assay need to be taken into account (Table 1). IL-8 was present in the supernatant of IL-1β-stimulated samples at concentrations around 50,000 pg/mL, whereas IL-6 and TNF-α were detected at 20- or even 1,000-fold lower concentrations. Stimulating for 16 h, Schuerer-Maly et al. (1994) reported similar IL-8 concentrations in HT-29, while in the supernatants of Caco-2 cells only 4 000 pg/mL were detected. Even though different exposure procedures render such direct comparisons difficult, the comparably high expression of IL-8 in our samples, leads to the assumption that some cellular signaling may be induced in HCEC-1CT cells substantially activating IL-8 expression and secretion. Thus, potential BUT-induced inhibiting effects on IL-8, might be overlaid by such co-occurring mechanisms.

Regarding cytotoxic effects, after 5 h of incubation the concentrations used in these experiments only marginally affected cellular viability. In the combinatory treatments cellular metabolic activity was slightly modulated, whereas no effects on the cellular protein content were observed. Various *in vitro* and *in vivo* studies investigated BUT-induced toxicities, showing the compound’s potential to trigger oxidative stress, to increase intracellular ROS production, lipid peroxidation and to cause oxidative DNA damage (Tookey et al., 1972; Burmeister et al., 1980; Vesonder et al., 1993; Wang et al., 2006; Liu et al., 2007; Wang et al., 2007; Shi et al., 2009; Wang et al., 2009a; Wang et al., 2009b; Yang et al., 2010). Whereas most previous studies administered comparably high concentrations of BUT, we could recently show in combinatory treatments with DON that BUT concentrations in the low micromolar range contributed significantly to the overall cytotoxicity of respective mixtures (Woelflingseder et al., 2018). Beyond that, pre-incubations with BUT for one or three hours were sufficient to enhance DON-induced cytotoxic effects (24 h) in HepG2 cells (Woelflingseder et al., 2018).

In conclusion, our results provide first evidences that the *Fusarium* secondary metabolite BUT, substantially alters the inflammatory response of the human intestinal epithelial cell line HCEC-1CT. NF-κB activity was significantly suppressed in the presence of BUT, resulting in downstream consequences such as reduced transcription and expression of pro-inflammatory cytokines. As BUT co-occurs in high abundance with mycotoxins such as DON and NX-3 (Abia et al., 2013; Streit et al., 2013b; Uhlig et al., 2013; Ivanova et al., 2016), known to act as activators of the NF-κB signaling pathway and to increase the expression of inflammatory genes in the intestine (Van De Walle et al., 2008; Alassane-Kpembi et al., 2017), we analyzed the intestinal inflammatory response following combined exposure to BUT and DON or NX-3. BUT, in 10-fold molar excess substantially suppressed the trichothecene-induced NF-κB activation, as well as the transcription and secretion of important pro-inflammatory cytokines. A relevant question is therefore whether such a situation may occur in naturally contaminated grain. A survey on 83 animal feed and feed components reported median concentrations of 23 and 122 *μ*g/kg of BUT and DON, respectively (Streit et al., 2013b). This suggests that on average the molar ratio BUT/DON is rather 0.4 than 10, and the observed interaction may not be very relevant. Yet, if individual samples are considered, a different picture emerges. In 3 of six corn cob samples, listed individually, the BUT/DON molar ratio was 29.2, 72.8 and 7.3, suggesting that an interference of BUT with DON-induced inflammatory responses could be relevant after all. Determination of BUT in mycotoxin surveys seems highly warranted to close the gap in occurrence data. In contrast to animal feed, grain used for food is typically not consumed raw, and the stability of BUT during food processing is also unknown.

Our observation of combinatorial effects between trichothecenes and BUT argue for further research into occurrence and intake of BUT, and a more detailed analysis of the underlying mechanism of action, in order to evaluate how BUT modulates the pathogenesis of intestinal inflammatory diseases and whether it should be considered in the risk assessment of naturally occurring mixtures.

## AUTHOR INFORMATION

### Author Contributions

LW was involved in designing the study, measuring the NF-κB reporter gene assay, qRT-PCR, cell viability and magnetic Luminex® assays, carried out statistical analysis, and wrote the manuscript. GA was involved in coordination of the project and refined the manuscript. DM was involved in designing and supervising the project and refined the manuscript.

## ACKNOWLEDGMENT

The authors would like to express their gratitude towards Nadia Gruber, performing preliminary NF-κB reporter gene assays, Julia Beisl for support during the performance of the Magnetic Luminex® assay and Maximilian Haider for the synthesis of butenolide.

## ABBREVIATIONS

ACTB1: β-actin
BUT: butenolide
COX-2: cyclooxygenase-2
d: days
DON: deoxynivalenol
GAPDH: glyceraldehyde 3-phosphate dehydrogenase
HKLM: heat killed Listeria monocytogenes
HO-1: heme oxygenase
IL-1α: interleukin-1α
IL-1β: interleukin-1β
IL-6: interleukin-6
IL-8: interleukin-8
iNOS: inducible nitric oxide synthase
JNK/SAPK: c-Jun N-terminal kinases/stress-activated kinases
LC-UV: liquid chromatography-ultraviolet
LPS: lipopolysaccharide
MAPKS: mitogen-activated protein kinases
MCP-1: monocyte chemotactic protein 1
NF-κB: nuclear factor kappalight-chain-enhancer of activated B cells
Nrf2/ARE: nuclear factor E2-related factor 2/antioxidant responsive element
qRT-PCR: quantitative reverse transcription polymerase chain reaction
ROS: reactive oxygen species
RT: room temperature
SRB: sulforhodamine B
TNF-α: tumor necrosis factor-alpha

## REFERENCES

Abia, W.A., Warth, B., Sulyok, M., Krska, R., Tchana, A.N., Njobeh, P.B., Dutton, M.F., and Moundipa, P.F. (2013). Determination of multi-mycotoxin occurrence in cereals, nuts and their products in Cameroon by liquid chromatography tandem mass spectrometry (LC-MS/MS). Food Control 31, 438–453.

Adesso, S., Autore, G., Quaroni, A., Popolo, A., Severino, L., and Marzocco, S. (2017). The Food Contaminants Nivalenol and Deoxynivalenol Induce Inflammation in Intestinal Epithelial Cells by Regulating Reactive Oxygen Species Release. Nutrients 9, 1343.

Alassane-Kpembi, I., Puel, O., Pinton, P., Cossalter, A.M., Chou, T.C., and Oswald, I.P. (2017). Co-exposure to low doses of the food contaminants deoxynivalenol and nivalenol has a synergistic inflammatory effect on intestinal explants. Arch Toxicol 91, 2677–2687.

Ali, Y., Alam, M.S., Hamid, H., Husain, A., Shafi, S., Dhulap, A., Hussain, F., Bano, S., Kharbanda, C., Nazreen, S., and Haider, S. (2015). Design and Synthesis of Butenolide-based Novel Benzyl Pyrrolones: Their TNF-alpha based Molecular Docking with In vivo and In vitro Anti-inflammatory Activity. Chem Biol Drug Des 86, 619–625.

Azcona-Olivera, J.I., Ouyang, Y., Murtha, J., Chu, F.S., and Pestka, J.J. (1995). Induction of cytokine mRNAs in mice after oral exposure to the trichothecene vomitoxin (deoxynivalenol): relationship to toxin distribution and protein synthesis inhibition. Toxicol Appl Pharmacol 133, 109–120.

Bennett, J.W., and Klich, M. (2003). Mycotoxins. Clin Microbiol Rev 16, 497–516.

Burkhardt, H.J., Lundin, R.E., and Mcfadden, W.H. (1968). Mycotoxins produced by Fusarium nivale (fries) cesati isolated from tall fescue (festuca arundinacea schreb.): Synthesis of 4-acetamido-4-hydroxy-2-butenoic acid-γ-lactone. Tetrahedron 24, 1225–1229.

Burmeister, H.R., Grove, M.D., and Kwolek, W.F. (1980). Moniliformin and butenolide: effect on mice of high-level, long-term oral intake. Appl Environ Microbiol 40, 1142–1144.

Dunne, A., and O’neill, L.A. (2003). The interleukin-1 receptor/Toll-like receptor superfamily: signal transduction during inflammation and host defense. Sci STKE 2003, re3.

Efsa, Knutsen, H.K., Alexander, J., Barregård, L., Bignami, M., Brüschweiler, B., Ceccatelli, S., Cottrill, B., Dinovi, M., Grasl-Kraupp, B., Hogstrand, C., Hoogenboom, L., Nebbia, C.S., Oswald, I.P., Petersen, A., Rose, M., Roudot, A.-C., Schwerdtle, T., Vleminckx, C., Vollmer, G., Wallace, H., De Saeger, S., Eriksen, G.S., Farmer, P., Fremy, J.-M., Gong, Y.Y., Meyer, K., Naegeli, H., Parent-Massin, D., Rietjens, I., Van Egmond, H., Altieri, A., Eskola, M., Gergelova, P., Ramos Bordajandi, L., Benkova, B., Dörr, B., Gkrillas, A., Gustavsson, N., Van Manen, M., and Edler, L. (2017). Risks to human and animal health related to the presence of deoxynivalenol and its acetylated and modified forms in food and feed. EFSA Journal 15, e04718–n/a.

Garreau De Loubresse, N., Prokhorova, I., Holtkamp, W., Rodnina, M.V., Yusupova, G., and Yusupov, M. (2014). Structural basis for the inhibition of the eukaryotic ribosome. Nature 513, 517–522.

Grove, M.D., Yates, S.G., Tallent, W.H., Ellis, J.J., Wolff, I.A., Kosuri, N.R., and Nichols, R.E. (1970). Mycotoxins produced by Fusarium tricinctum as possible causes of cattle disease. J Agric Food Chem 18, 734–736.

Hayden, M.S., and Ghosh, S. (2011). NF-kappaB in immunobiology. Cell Res 21, 223–244.

Iordanov, M.S., Pribnow, D., Magun, J.L., Dinh, T.H., Pearson, J.A., Chen, S.L., and Magun, B.E. (1997). Ribotoxic stress response: activation of the stress-activated protein kinase JNK1 by inhibitors of the peptidyl transferase reaction and by sequence-specific RNA damage to the alpha-sarcin/ricin loop in the 28S rRNA. Mol Cell Biol 17, 3373–3381.

Ivanova, L., Sahlstrøm, S., Rud, I., Uhlig, S., Fæste, C., Eriksen, G., and Divon, H. (2016). Effect of primary processing on the distribution of free and modified Fusarium mycotoxins in naturally contaminated oats. World Mycotoxin J 10, 1–16.

Jiang, T., Tian, F., Zheng, H., Whitman, S.A., Lin, Y., Zhang, Z., Zhang, N., and Zhang, D.D. (2014). Nrf2 suppresses lupus nephritis through inhibition of oxidative injury and the NF-kappaB-mediated inflammatory response. Kidney Int 85, 333–343.

Katika, M.R., Hendriksen, P.J., Van Loveren, H., and A, A.C.M.P. (2015). Characterization of the modes of action of deoxynivalenol (DON) in the human Jurkat T-cell line. J Immunotoxicol 12, 206–216.

Khokra, S.L., Monga, J., Husain, A., Vij, M., and Saini, R. (2013). Docking studies on butenolide derivatives as Cox-II inhibitors. Medicinal Chemistry Research 22, 5536–5544.

Kovalsky, P., Kos, G., Nahrer, K., Schwab, C., Jenkins, T., Schatzmayr, G., Sulyok, M., and Krska, R. (2016). Co-Occurrence of Regulated, Masked and Emerging Mycotoxins and Secondary Metabolites in Finished Feed and Maize-An Extensive Survey. Toxins 8, 363.

Laskin, J.D., Heck, D.E., and Laskin, D.L. (2002). The ribotoxic stress response as a potential mechanism for MAP kinase activation in xenobiotic toxicity. Toxicol Sci 69, 289–291.

Lin, W., Wu, R.T., Wu, T., Khor, T.O., Wang, H., and Kong, A.N. (2008). Sulforaphane suppressed LPS-induced inflammation in mouse peritoneal macrophages through Nrf2 dependent pathway. Biochem Pharmacol 76, 967–973.

Liu, J.B., Wang, Y.M., Peng, S.Q., Han, G., Dong, Y.S., Yang, H.Y., Yan, C.H., and Wang, G.Q. (2007). Toxic effects of Fusarium mycotoxin butenolide on rat myocardium and primary culture of cardiac myocytes. Toxicon 50, 357–364.

Liu, M., Zhou, Q., Wang, J., Liu, J., Qi, C., Lai, Y., Zhu, H., Xue, Y., Hu, Z., and Zhang, Y. (2018). Anti-inflammatory butenolide derivatives from the coral-derived fungus Aspergillus terreus and structure revisions of aspernolides D and G, butyrolactone VI and 4′,8′′-diacetoxy butyrolactone VI. RSC Advances 8, 13040–13047.

Liu, T., Zhang, L., Joo, D., and Sun, S.-C. (2017). NF-κB signaling in inflammation. Signal transduction and targeted therapy 2, 17023.

Lofgren, L., Riddle, J., Dong, Y., Kuhnem, P.R., Cummings, J.A., Del Ponte, E.M., Bergstrom, G.C., and Kistler, H.C. (2018). A high proportion of NX-2 genotype strains are found among Fusarium graminearum isolates from northeastern New York State. European Journal of Plant Pathology 150, 791–796.

Luo, Y., Eggler, A.L., Liu, D., Liu, G., Mesecar, A.D., and Van Breemen, R.B. (2007). Sites of alkylation of human Keap1 by natural chemoprevention agents. J Am Soc Mass Spectrom 18, 2226–2232.

Magesh, S., Chen, Y., and Hu, L. (2012). Small molecule modulators of Keap1-Nrf2-ARE pathway as potential preventive and therapeutic agents. Medicinal research reviews 32, 687–726.

Maresca, M., Yahi, N., Younes-Sakr, L., Boyron, M., Caporiccio, B., and Fantini, J. (2008). Both direct and indirect effects account for the pro-inflammatory activity of enteropathogenic mycotoxins on the human intestinal epithelium: stimulation of interleukin-8 secretion, potentiation of interleukin-1beta effect and increase in the transepithelial passage of commensal bacteria. Toxicol Appl Pharmacol 228, 84–92.

Moon, Y., Yang, H., and Lee, S.H. (2007). Modulation of early growth response gene 1 and interleukin-8 expression by ribotoxin deoxynivalenol (vomitoxin) via ERK1/2 in human epithelial intestine 407 cells. Biochem Biophys Res Commun 362, 256–262.

Nakamura, Y., and Miyoshi, N. (2010). Electrophiles in foods: the current status of isothiocyanates and their chemical biology. Biosci Biotechnol Biochem 74, 242–255.

Pestka, J.J., Zhou, H.R., Moon, Y., and Chung, Y.J. (2004). Cellular and molecular mechanisms for immune modulation by deoxynivalenol and other trichothecenes: unraveling a paradox. Toxicol Lett 153, 61–73.

Pinton, P., and Oswald, I.P. (2014). Effect of deoxynivalenol and other Type B trichothecenes on the intestine: a review. Toxins 6, 1615–1643.

Roig, A.I., Eskiocak, U., Hight, S.K., Kim, S.B., Delgado, O., Souza, R.F., Spechler, S.J., Wright, W.E., and Shay, J.W. (2010). Immortalized epithelial cells derived from human colon biopsies express stem cell markers and differentiate in vitro. Gastroenterology 138, 1012–1021.e1011-1015.

Roig, A.I., and Shay, J.W. (2010). Immortalization of adult human colonic epithelial cells extracted from normal tissues obtained via colonoscopy.

Schmittgen, T.D., and Livak, K.J. (2008). Analyzing real-time PCR data by the comparative C(T) method. Nat Protoc 3, 1101–1108.

Schuerer-Maly, C.C., Eckmann, L., Kagnoff, M.F., Falco, M.T., and Maly, F.E. (1994). Colonic epithelial cell lines as a source of interleukin-8: stimulation by inflammatory cytokines and bacterial lipopolysaccharide. Immunology 81, 85–91.

Shi, Z., Cao, J., Chen, J., Li, S., Zhang, Z., Yang, B., and Peng, S. (2009). Butenolide induced cytotoxicity by disturbing the prooxidant-antioxidant balance, and antioxidants partly quench in human chondrocytes. Toxicol In Vitro 23, 99–104.

Skehan, P., Storeng, R., Scudiero, D., Monks, A., Mcmahon, J., Vistica, D., Warren, J.T., Bokesch, H., Kenney, S., and Boyd, M.R. (1990). New colorimetric cytotoxicity assay for anticancer-drug screening. J Natl Cancer Inst 82, 1107–1112.

Soares, M.P., Seldon, M.P., Gregoire, I.P., Vassilevskaia, T., Berberat, P.O., Yu, J., Tsui, T.Y., and Bach, F.H. (2004). Heme oxygenase-1 modulates the expression of adhesion molecules associated with endothelial cell activation. J Immunol 172, 3553–3563.

Streit, E., Naehrer, K., Rodrigues, I., and Schatzmayr, G. (2013a). Mycotoxin occurrence in feed and feed raw materials worldwide: long-term analysis with special focus on Europe and Asia. J Sci Food Agric 93, 2892–2899.

Streit, E., Schwab, C., Sulyok, M., Naehrer, K., Krska, R., and Schatzmayr, G. (2013b). Multi-mycotoxin screening reveals the occurrence of 139 different secondary metabolites in feed and feed ingredients. Toxins 5, 504–523.

Sun, S.C., Chang, J.H., and Jin, J. (2013). Regulation of nuclear factor-kappaB in autoimmunity. Trends Immunol 34, 282–289.

Tookey, H.L., Yates, S.G., Ellis, J.J., Grove, M.D., and Nichols, R.E. (1972). Toxic effects of a butenolide mycotoxin and of Fusarium tricinctum cultures in cattle. J Am Vet Med Assoc 160, 1522–1526.

Ueno, Y. (1977). Mode of action of trichothecenes. Ann Nutr Aliment 31, 885–900.

Uhlig, S., Eriksen, G.S., Hofgaard, I.S., Krska, R., Beltran, E., and Sulyok, M. (2013). Faces of a changing climate: semi-quantitative multi-mycotoxin analysis of grain grown in exceptional climatic conditions in Norway. Toxins 5, 1682–1697.

Van De Walle, J., Romier, B., Larondelle, Y., and Schneider, Y.J. (2008). Influence of deoxynivalenol on NF-kappaB activation and IL-8 secretion in human intestinal Caco-2 cells. Toxicol Lett 177, 205–214.

Varga, E., Wiesenberger, G., Hametner, C., Ward, T.J., Dong, Y., Schofbeck, D., Mccormick, S., Broz, K., Stuckler, R., Schuhmacher, R., Krska, R., Kistler, H.C., Berthiller, F., and Adam, G. (2015). New tricks of an old enemy: isolates of Fusarium graminearum produce a type A trichothecene mycotoxin. Environ Microbiol 17, 2588–2600.

Varga, E., Wiesenberger, G., Woelflingseder, L., Twaruschek, K., Hametner, C., Vaclavikova, M., Malachova, A., Marko, D., Berthiller, F., and Adam, G. (2018). Less-toxic rearrangement products of NX-toxins are formed during storage and food processing. Toxicol Lett 284, 205–212.

Vesonder, R.F., Gasdorf, H., and Peterson, R.E. (1993). Comparison of the cytotoxicities of Fusarium metabolites and Alternaria metabolite AAL-toxin to cultured mammalian cell lines. Arch Environ Contam Toxicol 24, 473–477.

Wan, L.Y., Woo, C.S., Turner, P.C., Wan, J.M., and El-Nezami, H. (2013). Individual and combined effects of Fusarium toxins on the mRNA expression of pro-inflammatory cytokines in swine jejunal epithelial cells. Toxicol Lett 220, 238–246.

Wang, H.J., Wang, Y.M., and Peng, S.Q. (2009a). Repeated administration of a Fusarium mycotoxin butenolide to rats induces hepatic lipid peroxidation and antioxidant defense impairment. Food Chem Toxicol 47, 633–637.

Wang, Y.M., Peng, S.Q., Zhou, Q., Wang, M.W., Yan, C.H., Wang, G.Q., and Yang, H.Y. (2007). The oxidative damage of butenolide to isolated erythrocyte membranes. Toxicol In Vitro 21, 863–869.

Wang, Y.M., Peng, S.Q., Zhou, Q., Wang, M.W., Yan, C.H., Yang, H.Y., and Wang, G.Q. (2006). Depletion of intracellular glutathione mediates butenolide-induced cytotoxicity in HepG2 cells. Toxicol Lett 164, 231–238.

Wang, Y.M., Wang, H.J., and Peng, S.Q. (2009b). In ovo exposure of a Fusarium mycotoxin butenolide induces hepatic and renal oxidative damage in chick embryos, and antioxidants provide protections. Toxicol In Vitro 23, 1354–1359.

Wardyn, Joanna d., Ponsford, Amy h., and Sanderson, Christopher m. (2015). Dissecting molecular cross-talk between Nrf2 and NF-κB response pathways. Biochemical Society Transactions 43, 621–626.

Warhurst, A.C., Hopkins, S.J., and Warhurst, G. (1998). Interferon gamma induces differential upregulation of alpha and beta chemokine secretion in colonic epithelial cell lines. Gut 42, 208–213.

Weber, A., Wasiliew, P., and Kracht, M. (2010). Interleukin-1 (IL-1) pathway. Sci Signal 3, cm1.

Woelflingseder, L., Del Favero, G., Blazevic, T., Heiss, E.H., Haider, M., Warth, B., Adam, G., and Marko, D. (2018). Impact of glutathione modulation on the toxicity of the Fusarium mycotoxins deoxynivalenol (DON), NX-3 and butenolide in human liver cells. Toxicol Lett 299, 104–117.

Yang, H.Y., Wang, Y.M., and Peng, S.Q. (2010). Metallothionein-I/II null cardiomyocytes are sensitive to Fusarium mycotoxin butenolide-induced cytotoxicity and oxidative DNA damage. Toxicon 55, 1291–1296.

Yates, S.G. (1971). Toxin-Producing Fungi from Fescue Pasture. New York: Academic Press.

Yates, S.G., Tookey, H.L., Ellis, J.J., Tallent, W.H., and Wolff, I.A. (1969). Mycotoxins as a possible cause of fescue toxicity. Journal of Agricultural and Food Chemistry 17, 437–442.

Zhou, H.R., Yan, D., and Pestka, J.J. (1997). Differential cytokine mRNA expression in mice after oral exposure to the trichothecene vomitoxin (deoxynivalenol): dose response and time course. Toxicol Appl Pharmacol 144, 294–305

